# Precise measurements of chromatin diffusion dynamics by modeling using Gaussian processes

**DOI:** 10.1101/2021.03.16.435699

**Authors:** Guilherme M. Oliveira, Attila Oravecz, Dominique Kobi, Manon Maroquenne, Kerstin Bystricky, Tom Sexton, Nacho Molina

## Abstract

The spatiotemporal organization of chromatin influences many nuclear processes: from chromosome segregation to transcriptional regulation. To get a deeper understanding of these processes it is essential to go beyond static viewpoints of chromosome structures, and to accurately characterize chromatin mobility and its diffusion properties. Here, we present GP-FBM: a new computational framework based on Gaussian processes and fractional Brownian motion to analyze and extract diffusion properties from stochastic trajectories of labeled chromatin loci. GP-FBM is able to optimally use the higher-order correlations present in the data and therefore outperforms existing methods. Furthermore, GP-FBM is able to extrapolate trajectories from missing data and account for substrate movement automatically. Using our method we show that diffusive chromatin diffusion properties are surprisingly similar in interphase and mitosis in mouse embryonic stem cells. Moreover, we observe surprising heterogeneity in local chromatin dynamics, which correlates with transcriptional activity. We also present GP-Tool, a user-friendly graphical interface to facilitate the use of GP-FBM by the research community for future studies of nuclear dynamics.

## Introduction

The spatiotemporal organization of chromatin plays a crucial role in several nuclear processes: from cell division, where chromatin is compacted during mitosis facilitating chromosome segregation, to gene regulation, where precise control of transcription correlates with specific long-range chromatin contacts [1]. Chromosome conformation capture techniques and imaging approaches have revealed fundamental structural features of chromatin at different resolutions. Of special interest are topological associated domains (TADs) which are characterized by an increased frequency of interactions between genomic loci within the same domain, and reduced interactions across domains [2, 3]. Remarkably, it has been shown that TAD organization can influence regulation of transcription [4] and that TADs are dismantled during mitosis where chromatin density dramatically increases [5, 6]. However, much less is known about the diffusion and mobility properties of chromatin and how they depend on the genomic context. For instance, it is not clear whether or how transcriptional activation affects chromatin mobility, with previous studies giving seemingly conflicting results [7, 8]. Insights into the dynamic properties of chromatin motion are required to understand how gene regulatory elements explore the nuclear space to “find “ their distal targets and/or to build regulatory hubs [9].

The simplest model to describe the diffusion of microscopic systems is Brownian motion, whereby movements are caused by random collisions of small particles within the system [10, 11]. However, as a bulky polymer making interactions with the nuclear environment, chromatin frequently displays sub-diffusive behavior, more constrained than classical Brownian motion [12]. Therefore, the mean squared displacement (MSD) of chromatin is expected to follow this relationship with time: *MSD* ∝ *Dt*^*α*^. Two parameters thus describe the diffusion properties of chromatin: the apparent diffusion coefficient *D*, indicating the speed of motion, and the anomalous diffusion exponent *α*, which for sub-diffusive behavior is *<* 1 which indicates greater confinement (henceforth we refer to it as confinement parameter). The traditional method used to estimate the diffusion parameters is based on calculating the MSD over time from measured trajectories and fitting the above theoretical expression to the data. More sophisticated methods, such as can be derived from Spot-On [13], use the whole distribution of observed displacements (henceforth referred to as DDB (displacement distribution-based) methods). However, they do not use all the information contained in the trajectories as higher-order correlations are discarded. Furthermore, errors due to measurement noise cannot easily be included into the analysis, and it is not possible to recover missing data points due to misdetection or occlusions.

Here, we propose a novel computational method, GP-FBM, based on Gaussian Processes (GP) [14] and fractional Brownian motion (FBM) [15, 16]. Importantly, GP provides a consistent probabilistic framework that considers entire trajectories and thus utilizes all the available information. Trials on simulated data demonstrate a greater precision of GP-FBM in measuring diffusion parameters over existing techniques. Furthermore, GP-FBM naturally takes into account imprecision in localization and occlusions without the need to establish a fitted MSD curve or displacement distributions, as it is applied directly on trajectories. We further extend this model to account for external sources of movement (e.g. displacement of whole nuclei or chromosomes) using underlying correlations between multiple trajectories, without the need to further develop substrate motion models and experiments for calibration. Finally, we applied GP-FBM to two experimental systems to study chromatin diffusion properties in different contexts. First, we characterized chromatin dynamics in interphase and mitosis using tagged arrays inserted at random genomic locations in mouse embryonic stem (ES) cells. Although chromatin density increases by a factor of three during mitosis [17], our results surprisingly indicate that there are no significant differences on average in the diffusion coefficient or confinement parameter. Second, to compare the diffusion properties of different specific genomic regions, we performed double-labeling and live tracking experiments around the HoxA locus in mouse ES cells before and after induction of the genes with retinoic acid. We discover that, instead of having homogeneous diffusion properties across euchromatin, genomic loci significantly differ in both their apparent diffusion coefficients and confinement parameters. In some cases, altered chromatin diffusion properties correlate with underlying functions such as gene activity changes. The methods we have developed have been integrated into a user-friendly package, GP-Tool, for use in the scientific community. Chromatin mobility has been largely overlooked in previous studies of genome functions, and we anticipate that GP-FBM will greatly facilitate research in that area.

## Results

### GP-FBM: A Gaussian process with a fractional Brownian kernel to model particle diffusion dynamics

Current methods to analyze particle diffusion dynamics rely on two-point statistics as only particle displacements calculated between two frames at different time intervals are used. This has important drawbacks: higher order correlations within the trajectories are discarded; errors due to measurement noise cannot easily be included into the analysis; and missing data points due to misdetection or occlusions are ignored. To address these problems we built a consistent probabilistic framework based on Gaussian Processes (GP). Briefly, a GP is defined as a collection of random variables such that every finite subset of them follows a multivariate normal distribution which is fully determined by its mean and kernel functions *µ*(*t*) and Σ(*t, t*^*i*^) [14]. We assume that a stochastic diffusion trajectory *x*(*t*) of a given chromatin locus can be modeled as a Gaussian process with the following fractional Brownian kernel:

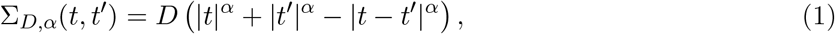

where *D* is the apparent diffusion coefficient and *α* is the confinement parameter defined in the range 0 *< α <* 2. This kernel produces a generalized Brownian motion with a mean squared displacement (*r*^2^) = 2*nDt*^*α*^, where *n* corresponds to the number of degrees of freedom. Then the probability of observing a discrete trajectory **r** = {*r*_*i*_} measured at a set of times ***t*** = {*t*_*i*_} is given by the multivariate Gaussian distribution,

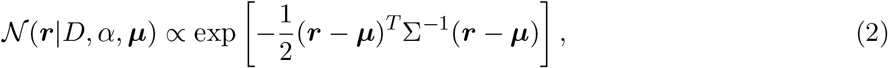

where the covariance matrix is defined as Σ_*ij*_ ≡ Σ_*D,α*_(*t*_*i*_, *t*_*j*_) and we take a constant ***µ*** without loss of generality. Furthermore, we can easily incorporate localization errors by adding the diagonal term 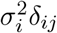 to the covariance matrix Σ, which assumes that errors are decorrelated and normally distributed with standard deviations ***σ*** = {*σ*_*i*_} (see Methods). Ultimately, providing a trajectory ***r*** and the localization errors ***σ***, the likelihood (2) can be used to calculate estimates of the diffusion parameters *D* and *α*.

To test the performance of the GP-FBM method, we generated 2000 synthetic trajectories using uniformly distributed random values of *D* and *α* in the range 0 *< D <* 1.5 and 0 *< α <* 2. As real trajectories present occlusions and localization imprecision, we removed 10% of the points and introduced positional noise of about 1*/*10 of a pixel (an example is shown in Fig. 1a). We then obtain a posterior distribution over the parameters *D* and *α* given the trajectory and the localization errors by combining the likelihood (2) with flat priors over the parameters, which can be sampled using Markov Chain Monte Carlo (MCMC) (see Fig. 1b and Methods). Interestingly, once the diffusion parameters are estimated, the power of the GP framework can be used to infer the most probable trajectory of the particle by removing measurement noise and predicting the localization of the particle position where occlusions or misdetections occurred (Fig. 1a). We compared the results obtained using our approach on the simulated data with the traditional MSD method and a DDB method inspired by Spot-On [13]. GP-FBM clearly outperforms both methods, producing much smaller relative errors of parameter estimation (Fig. 1c). Unlike GP-FBM, both MSD and DDB methods require trajectories to be split into individual displacements, thus breaking higher order correlations that the trajectories may contain. Therefore GP-FBM method can optimally infer diffusion parameters from single trajectories, using all the information contained in the data and thus achieving greater precision.

**Figure 1.**
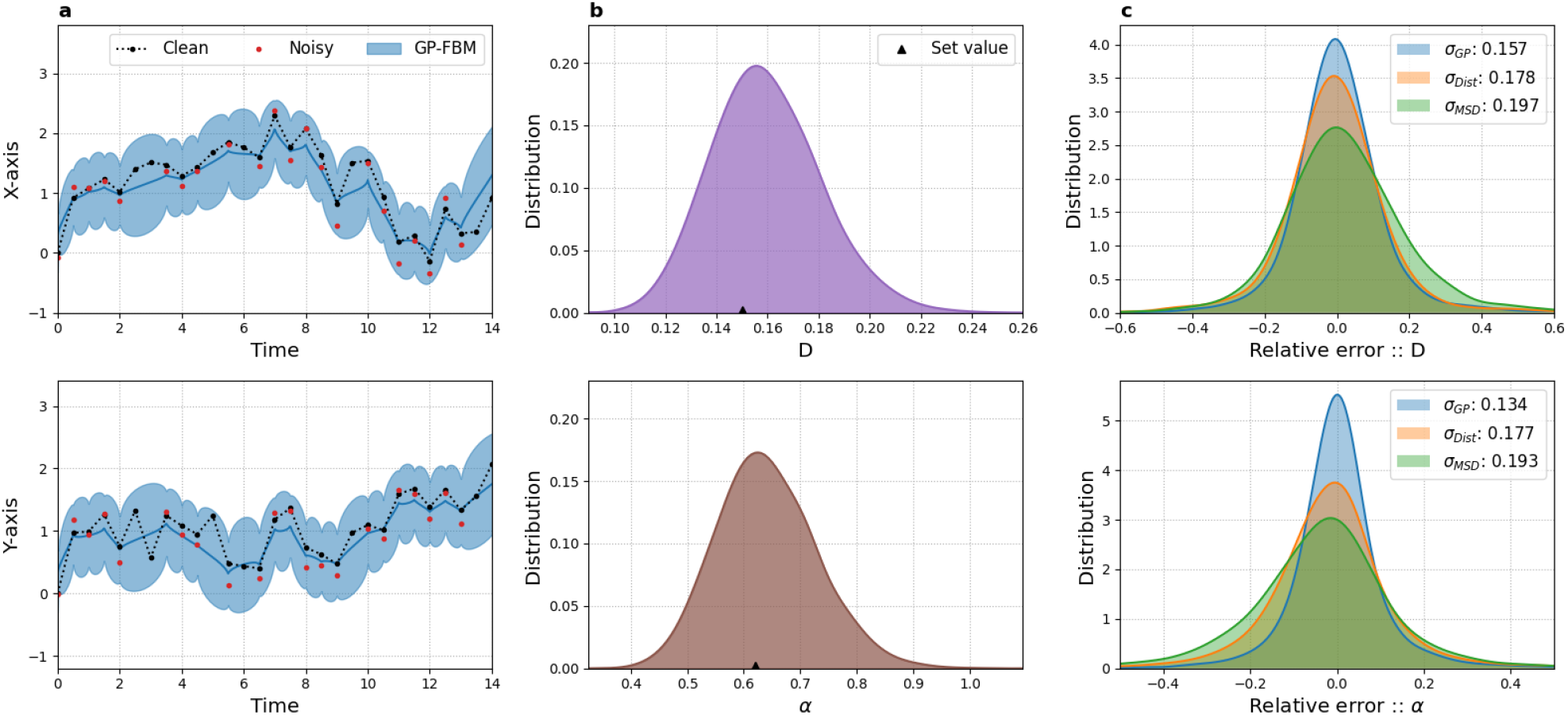
GP-FBM outperforms existing methods on simulated data. (a) Simulated 2D trajectory and GP-FBM inference: black dots are simulated values, red dots are the values after inclusion of noise and 10% occlusions, while the blue line is the most probable trajectory with shading representing the 95% credible interval as inferred by GP-FBM. (b) Posterior probability distributions for *D* (top) and *α* (bottom) inferred for trajectory in (a). Triangles denote the values set in the simulation. (c) Comparing inference relative errors as obtained by GP-FBM (blue), MSD (green) and DDB (orange) for 2000 simulated trajectories with random diffusion parameters.

### Accounting for substrate movement with GP-FBM

Often a particle may be subject to secondary movement that is entangled with its diffusion dynamics. In chromatin dynamics, this movement is frequently associated with the substrate in which the particle is diffusing, such as cell displacement, membrane fluctuations or chromatin reallocation, as well as technical considerations such as thermal drift and undesired media flow. If overlooked, this may result in over-estimation of the diffusive properties. However, when two or more particles are measured in the same cell, this substrate movement can be accounted for by analyzing the correlation that introduces between the particle trajectories. To that end, we developed a correlation covariance model that takes advantage of the GP-FBM framework to quantify and handle the substrate movement and the possible correlations that may introduce into the movement of particles (see Methods). In the case of two particles, we obtain the probability distribution,

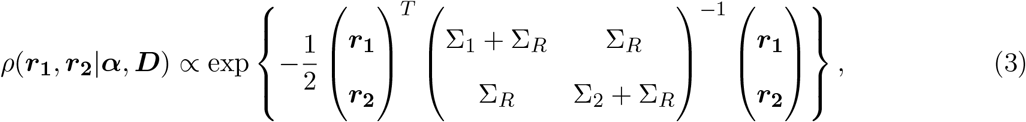

where Σ_1_, Σ_2_ and Σ_*R*_ are FBM covariance matrices for the two particles and substrate respectively with diffusion parameters ***D*** = {*D*_1_, *D*_2_, *D*_*R*_} and ***α*** = {*α*_1_, *α*_2_, *α*_*R*_}. This method can easily be extended for higher numbers of particles, limited only by required computational power in practice. In this study, we restrict our analysis to five particles per cell.

To demonstrate the utility of this approach, we generated 2000 synthetic trajectories as before, but now including substrate movement, generating vectors ***r***_*i*_ as a combination of substrate displacement ***R*** and the actual particle displacement ***a***_*i*_ (Fig. 2a). Again, simulations are generated with 10% occlusion rate and localization errors. As expected, without applying the correlation covariance model, *D* and *α* tend to be overestimated; however, the parameters are precisely measured when the substrate correction is incorporated into the model (Fig. 2b,c). The method is also able to precisely estimate the dynamic properties of the substrate and to estimate, albeit with less precision, the substrate movement (see Fig. 2d, Fig. S1 and Methods). Furthermore, we tested the performance of the method depending on the number of tracked particles subjected to the same substrate movement. Precision is increased with use of more particles, but the bulk of the error is already removed with only two particles (see Fig. S2). Finally, we showed that GP-FBM outperforms a DDB method where the substrate movement is taken into account (see Methods and S3). In conclusion, GP-FBM has the ability to remove the substrate movement from the analysis which is demonstrably important for precise measurement of diffusion parameters. Other approaches can be applied to estimate cell movements and correct trajectories [18], but GP-FBM has the advantage of being able to derive this information directly from the trajectories themselves, provided that two or more particles are tracked per cell, thanks to the correlation that the substrate movement introduces between the particles.

**Figure 2.**
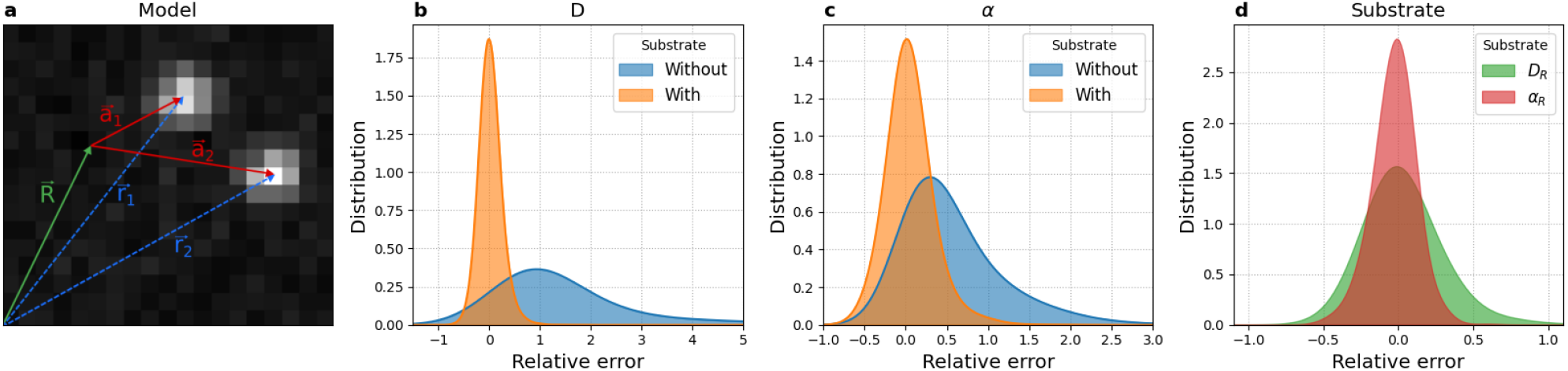
GP-FBM can remove substrate movement to improve estimation of diffusion parameters. (a) Scheme showing the measured trajectories of the particles ***r***_**1**_ and ***r***_**2**_ as the combination of substrate motion ***R*** and the particles ‘ diffusion with respect to the substrate, ***a***_1_ and ***a***_2_. (b-c) Distributions of relative errors for the estimation of diffusion parameters on 2000 pairs of simulated trajectories with random parameters when substrate movement is accounted for (orange) or not (blue). (d) Distributions of relative errors for the estimation of diffusion parameters of substrate motion.

### Analyzing chromatin dynamics in interphase and mitosis with GP-FBM

Due to chromosome compaction and condensation, chromatin density increases by a factor of three during mitosis [17]. Although, the structure of mitotic chromatin has been intensively studied [5, 19, 20], it is unknown if or how the higher density and rearrangement of chromatin fibers affects chromatin diffusion properties. To measure chromatin dynamics in interphase and mitosis, we used a mouse ES cell line carrying approximately 20 *TetO* arrays of 7 kb length inserted at random genomic locations [21]. GFP::TetR is stably expressed in these cells, where it binds to the *TetO* arrays for the simultaneous visualization of several chromatin loci in each cell. We performed confocal live-imaging of these cells, distinguishing interphase and mitotic cells by DNA staining using Hoechst 33342, recording images at 4 frames/s for 75 s. To increase the number of mitotic cells we also performed live-imaging experiments on cells arrested in prometaphase with nocodazole (see Fig 3a and Methods). Before applying the GP-FBM probabilistic framework, we enhanced particle localization precision by fitting a 2D Gaussian function to the intensity of the tracked spots (see Methods and S4). Comparing the performance of GP-FBM with and without substrate movement correction, it was apparent that actively dividing mitotic cells had greater substrate movement (presumably due to coordinated alignment and movement of chromosomes by the mitotic spindle), but that appreciable correction was required for precise chromatin dynamics measurements in all conditions (Fig. 3b). Surprisingly, we observed no significant differences in the mean diffusion coefficient or confinement parameter between interphase and mitotic chromosomes, suggesting that condensation may not necessarily affect the average local diffusion dynamics of chromatin (Fig. 3c and d). We observed a small but significant increase in the confinement parameter of mitotic-arrested cells compared to interphase, which might be related to the effect that nocodazole has on microtubule formation and thus mitotic chromosome stability [22].

**Figure 3.**
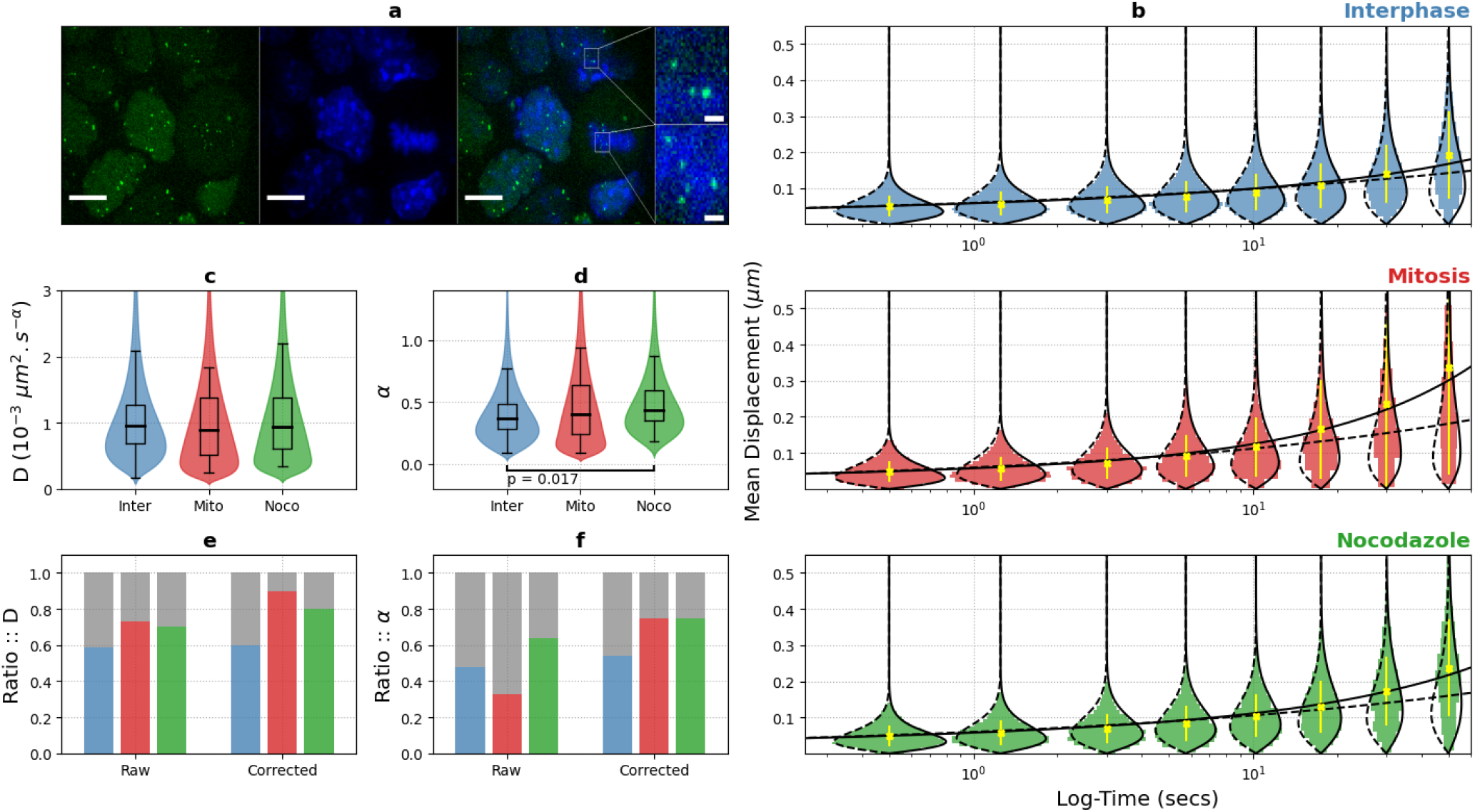
Chromatin dynamics are similar in interphase and mitosis, and are highly variable across loci. (a) Maximum projection images of ES cells containing spots of TetR::GFP (green) bound to *TetO* arrays, with DNA (blue) stained with Hoechst (scale bar is 10 *µm*). Inset shows magnification of the selected area (scale bar 1 *µm*). (b) Displacement distributions for loci in interphase (blue), mitosis (red) or mitotic arrest (green), plotted for different time intervals. The observed mean displacement is shown with yellow crosses and theoretical curves with and without substrate movement are shown with black continuous and dashed lines, respectively. The model with substrate movement fits better the data specially at greater time intervals, suggesting a greater confounding effect, especially for active mitotic cells. (c-d) Distribution of estimated *D* and *α* in the three conditions, correcting for substrate movement. (e-f) Estimations of the inter-(grey) and intra-cell (color) proportions of total variance for *D* and *α* in interphase (blue), mitosis (red) and mitotic arrest (green).

Interestingly, we obtained a wide range of estimated *D* and *α* coefficients indicating a remarkable spot-to-spot variability in their diffusion dynamics, even when correcting for substrate movement. This variability could partially be caused by differences in the state of the analyzed cells (inter-cell variability) leading to different overall chromatin dynamics. Alternatively, differences in the chromatin context of the genomic loci could lead to specific diffusion dynamics (intra-cell variability). Applying the law of total variance, we quantified the contribution of inter-cell vs intra-cell variability (see Methods) (Fig. 3e and f). Strikingly, as much as 75% of the variability in *D* and 65% in *α* could be explained by differences *within* the same cells, and this estimate is even higher when substrate movement is taken into account. Together, this suggests that different genomic loci may have characteristic diffusion dynamic properties due to their specific chromatin or nuclear context.

### GP-FBM distinguishes locus- and cell-specific chromatin diffusion properties

Except for a tendency for chromatin mobility to be reduced at centromeric or telomeric locations in yeast [23], little is known about how different genomic contexts may affect dynamics of the underlying chromatin. Further, previous studies give conflicting views on whether transcriptional activation can increase local confinement of a gene (as observed in the same cell before and after estrogen stimulation [24]) and/or increase gene mobility (as observed comparing cells before and after differentiation [25]). To compare the diffusion properties of different specific genomic regions, we performed double-labeling and live tracking experiments with the ANCHOR system [26] around the HoxA locus in mouse ES cells before and after induction of Hox genes with retinoic acid. We engineered the ANCH1 and ANCH3 [24] labels into different locations within the same allele to generate two ES lines with equidistant probes assessing inter-TAD (T1-T2) or intra-TAD (T2-T3) associations (Fig. 4a), and imaged at 2 frames/s for 2 min. As may be expected, the average inter-probe distance was higher for the inter-TAD than intra-TAD combination, but with large heterogeneity in the distance distributions (Fig. 4b; [27]). Interestingly, Hox gene induction had no effect on intra-TAD distances within the neighboring domain, but increased inter-TAD distances, supporting the idea of general TAD reinforcement as cell differentiation is induced [28]. We performed GP-FBM for the three loci and found that, in undifferentiated ES cells, although they have equivalent diffusion rates, region T1 is significantly more confined than T2 or T3 (Fig. 4c,d). Closer inspection of the ES (and differentiated neuronal precursor cell) epigenomic profiles around these regions showed that T1 is close (<15 kb) to a putative active enhancer of *Halr1* (Fig. S5). This gene encodes the long non-coding RNA *Haunt*, whose specific expression in ES cells is linked to suppression of the HoxA genes [29]. Active histone modifications around T1, compared to the silent T2 and T3 regions, thus appears linked to a greater confinement of the chromatin, in line with a previous study of an estrogen-induced gene [24]. Hox gene induction by retinoic acid had no significant effect on the diffusion rate of T1 but did reduce locus confinement (Fig. 4c,d). In contrast, the region T2, which lacks any known epigenomic or regulatory features, had increases in D and *α*, perhaps indicative of general chromatin remodeling caused on onset of differentiation. Curiously, T3 became more confined on retinoic acid treatment, with a concomitant increase in diffusion rate. This region contains sites bound by the architectural protein CTCF, whose binding is either lost or reduced on differentiation to neuronal precursors (Fig. S5). CTCF is proposed to form a roadblock for cohesin-mediated loop extrusion processes [30, 31], and this may be expected to play out in alterations to local chromatin dynamics, although this has been largely unexplored. Overall, these results show a correlation between activity or function of a gene locus and its local chromatin dynamics, a feature that has been largely overlooked in most previous studies.

**Figure 4.**
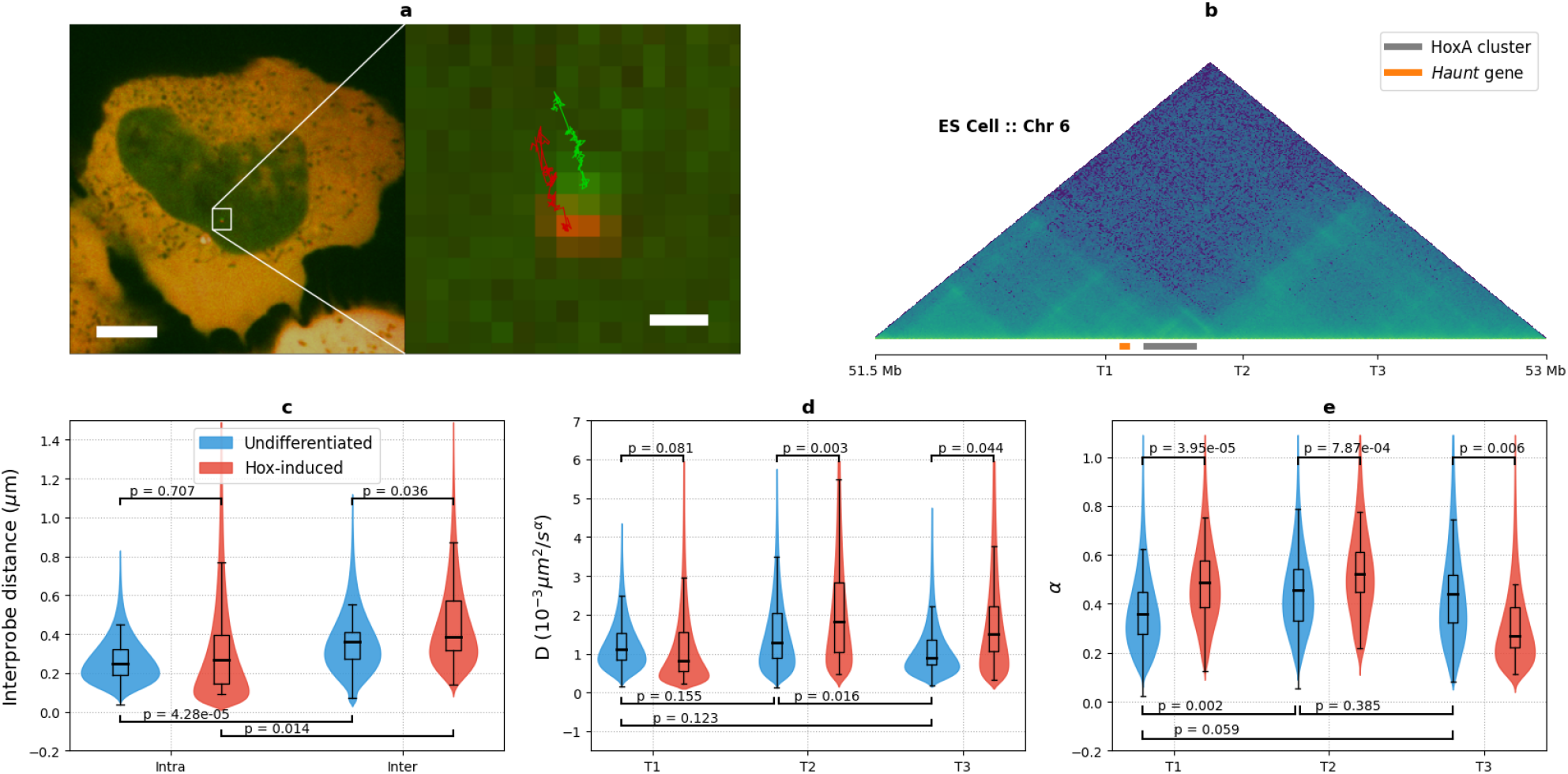
Chromatin dynamics of three chromatin loci within the HoxA genomic region. (a) Example of microscopy image obtained on ANCHOR cell line (scale 5 *µm*) with tracked trajectories shown in amplified subpanel (scale 0.3 *µm*). (b) ES Hi-C map around the HoxA cluster, illustrating the locus TAD structure. The positions of ANCHOR probes (T1, T2 and T3) and the genes *Haunt* and the HoxA cluster are shown underneath the map. (c) Inter-probe distances measured for ES cells before (blue) and after (red) treatment with retinoic acid. (d-e) Comparisons of diffusion coefficients and confinement parameters between the three labeled loci, before (blue) and after (red) treatment with retinoic acid.

### GP-Tool: A user-friendly graphical interface to apply GP-FBM

To facilitate use of GP-FBM by the community, we developed a freely available graphical user interface called GP-Tool (Fig. 5; github.com/guilmont). GP-Tool contains 4 plugins: movie, alignment, trajectories and g-process. The movie plugin allows the user to open TIFF files, display basic ImageJ and OME metadata, define colormaps for each channel and manually correct for contrast. The alignment plugin runs the algorithm described in Methods to digitally correct chromatin aberration and possible camera calibration issues. Alternatively, the user can manually modify each of the parameters. The program can perform the analysis of several cells in the same movie. The trajectories plugin provides the ROI utility which allows the user to select spots of interest and extract trajectories. Finally, the g-process plugin allows to infer the diffusion and confinement parameters correcting for substrate movement if two or more particles are tracked. It is also possible to use a MCMC sampler to obtain the posterior probability distribution associated with each of these parameters. Once the analysis is complete, the tool provides the possibility to save the results into two file formats: JSON and HDF5.

**Figure 5.**
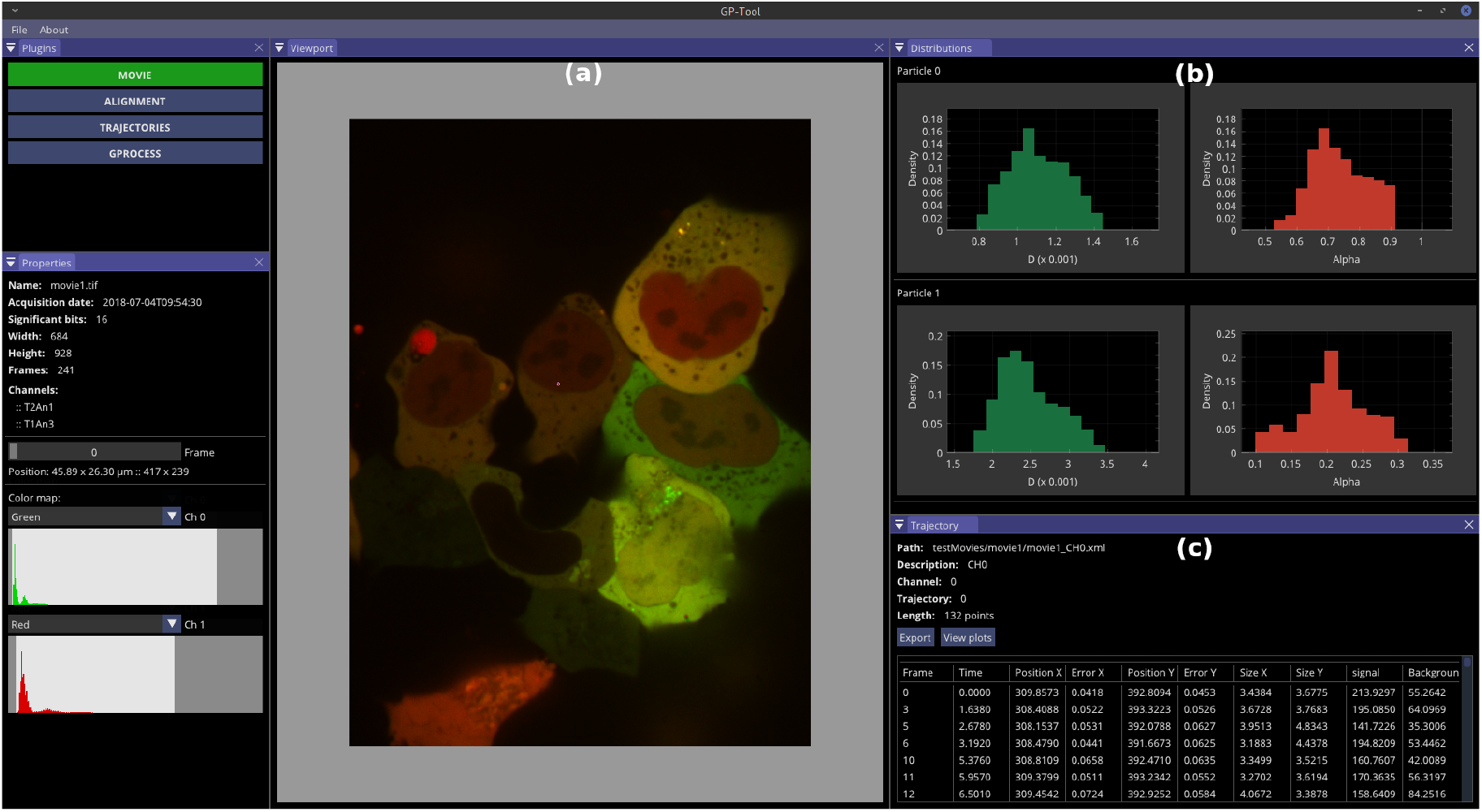
GP-Tool program: A graphical user interphase to apply GP-FBM on microscopy movies. (a) Main viewport (b) GProcess plugin also calculates the posterior distribution for *D* and *α* using MCMC. (c) The plugin for trajectories enhances the localization of tracked trajectories, estimate error, particle size and signal.

It also provides export functions to save tables in CSV format. All these formats are easily parsed in all major computing languages, such as C/C++, Python and R.

## Discussion

We developed GP-FBM, a Bayesian framework that combines the inference power of Gaussian processes with fractional Brownian motion, a flexible model to describe diffusion dynamics. GP-FBM treats stochastic trajectories as a whole without preprocessing or extracting limited statistics from them. Therefore, this approach utilizes optimally all the information contained in the data by incorporating higher-order correlations into the analysis. Furthermore, the Gaussian process framework allows us to easily integrate spot-dependent localization errors which translates into a consistent weighting of the time point depending on the precision at which the spot position is determined. Furthermore, missing data due to spot misdetection or occlusion does not hinder the analysis and, on the contrary, GP-FBM can be used to infer spot positions at time points when it was not measured. Finally, when two or more particles are tracked inside the cell, GP-FBM uses possible cross-correlation between the trajectories to characterize substrate movement and, therefore, remove it from the analysis of the diffusion dynamics of particles. Benchmarking shows improved precision of GP-FBM against two existing methods: the traditional MSD curve fitting and the more advanced DDB approach. This increase in accuracy can potentially be crucial to study changes in diffusion dynamic properties in different conditions, and can even be applied to single trajectories.

We applied GP-FBM to two ES cell systems to ask different questions about chromatin diffusion properties. Overall, we observe a large variability in chromatin dynamics when comparing individual cells and comparing different loci. A fraction of this variability can be explained by differences across cells, especially in interphase cells, indicating that cell state related to cell cycle or metabolism may globally influence chromatin dynamics. However, the majority of the observed variability is related to differences across loci. On the other hand, chromatin exhibits on average similar diffusion dynamics in interphase and mitosis despite a large difference in chromatin density. This result may be related with recent findings showing that mitotic chromatin is not as inaccessible and inert as previously thought. Indeed, several studies have shown that mitotic chromatin is bound by transcription factors [32, 33] and some genes are even transcribed during mitosis [34]. In contrast, different genomic loci can have striking differences from average chromatin dynamic properties, which appear to correlate with underlying functions such as transcriptional activity or binding of insulator proteins. More widespread application of GP-FBM to labeled transcribed loci and other specific regulatory elements, such as enhancers or TAD borders, will likely uncover more interesting functional links between genome function and the dynamics of its component chromatin.

Finally, we present GP-Tool a graphical user interface that helps to perform GP-FBM analysis on microscopy movies with only a few mouse clicks. Importantly, this tool and the GP-FBM framework can be applied to study not only chromatin dynamics but potentially any labeled particle that can be tracked over time. Moreover, the FBM kernel used in this study can easily be replaced in principle by alternative kernels that may better describe dynamics of other systems. We thus anticipate that GP-FBM and GP-Tool will greatly facilitate the analysis of diffusion dynamics in biology.

## Methods

### Cell lines, culture and treatments

#### Transgenic TetO ES line

The mouse ES cell line was kindly provided by Dr. Luca Giorgetti. It is derived from an X0 clone of the PGKT2 subclone of the feeder-independent PGK12.1 mouse ES line which was engineered by co-transfection with pBROAD3-TetR-ICP22NLS-eGFP and pcDNA3.1Hygro to stably express the TetR-eGFP recombinant protein after random integration and hygromycin selection (250 *µ*g/ml) as described in [21, 35]. The piggyBac transposon system was then used to generate cells with 20-25 stable random integrations of a 150 TetO binding site array as described in [21]. Cells were cultured on 0.1% gelatin-coated culture plates in DMEM (4.5g/l glucose) supplemented with GLUTAMAX-I, 15% fetal calf serum (ES cell culture tested), 0.1 mM beta-mercaptoethanol, 1,500 U/ml leukemia inhibitory factor (LIF; produced in house) and 0.1 mM non-essential amino acids in 5% CO_2_ at 37*°* C. Mitotic arrest was performed by treating the cells for 5 h with 100 ng/ml Nocodazole (Sigma, M1404-2MG).

#### Transgenic ANCHOR ES lines

J1 mouse ES cells were grown on gamma-irradiated mouse embryonic fibroblast cells under standard conditions (4.5 g/L glucose-DMEN, 15% FCS, 0.1 mM non-essential amino acids, 0.1 mM beta-mercaptoethanol, 1 mM glutamine, 500 U/mL LIF, gentamicin), then passaged onto feeder-free 0.2% gelatin-coated plates for at least two passages to remove feeder cells before subsequent transfections. The two (“inter-TAD “ and “intra-TAD “) ANCHOR transgenic lines were generated by sequential CRISPR/Cas9-mediated knock-in experiments in the following manner. First, flanking homology arms (mm9 chr6: 52,320,061-52,321,144, and chr6: 52,321,145-52,322,244) were introduced by PCR amplification and Gibson assembly into a vector containing ANCH1 sequence [26]. This vector (1 *µ* g) was co-transfected with 3 *µ*g of a vector containing Cas9-GFP, a puromycin resistance marker and the scaffold to transcribe the sgRNA specific to the T2 insertion site (CGGCGCGCACTTAACACCAA; vector generated by the IGBMC Molecular Biology platform) in 1 million cells with Lipofectamine-2000. Two days after transfection, the cells were cultured for 24 h with 3 *µ*g/ml puromycin, then 48 h with 1 *µ*g/ml puromycin to enrich for transfected cells, before sorting individual GFP-positive cells on to feeders to amplify individual clones. Clones with the correct sequence were screened by PCR and sequencing, then the CRISPR knock-in process was repeated to insert the ANCH3 sequence [24] into either the T1 site (“inter-TAD “ line; homology arms at chr6: 52,013,471-52,014,370 and chr6: 52,014,371-52,015,270; gRNA sequence AATCGAGCTCACGCCATTAG) or the T3 site (“intra-TAD “ line; homology arms at chr6: 52,622,955-52,623,855 and chr6: 52,623,856-52,624,755; gRNA sequence TATGCTGAGGCGTGTCGCAA). Final clones were verified for maintained pluripotency by qRT-PCR to assess Oct4, Nanog (e.g. Fig. S5) and Sox2 expression. Subsequent microscopy experiments (see below) confirmed heterozygous incorporation of the ANCH sequences (detection of one specific spot per ANCH sequence per cell) within the same allele (two spots were always in close proximity).

#### OR transfection

150,000 cells are plated two days prior to imaging off feeder cells onto laminin-511-coated 35 mm glass bottom petri dishes, and transfected with 3 *µ*g OR1-EGFP and 3 *µ*g OR3-IRFP plasmids (vectors available from NeoVirTech; were modified from original source by changing the C-terminal fluorescent protein sequence, introducing Kozak sequence before the translation start site and replacing the CMV promoter with EF-1*α*) using Lipofectamine-2000. After two days, the medium is changed to remove dead cells, before passing directly to microscopy.

#### Hox induction

ES cells were passaged without feeders and cultured on laminin-511 for two days without LIF, then for a subsequent three days without LIF and with the addition of 5 *µ*M retinoic acid. One day after the addition of retinoic acid, the cells are transfected with the OR proteins as previously.

### Microscopy

#### Live cell imaging of TetO ES cells

35 mm glass-bottom dishes (Ibidi 81158) were coated with 10 *µ*g/ml fibronectin human plasma (Sigma, F2006-1MG) in PBS for 45 minutes at room temperature. 3-5×105 cells were seeded one day before imaging, then the medium was replaced by phenol-red-free medium containing 500 ng/ml Hoechst 33342 (Invitrogen, H3570). Cells arrested in mitosis were collected on the day of imaging by “shake- off “, incubated with 0.25% Trypsin-1 mM EDTA (Invitrogen, 25200-072) for 1 min at 37°C and washed, and placed on fibronectin-coated glass-bottom dishes in phenol-red-free medium containing 100 ng/ml Nocodazole and 500 ng/ml Hoechst 33342. Confocal live-cell imaging was performed on a Nikon Eclipse Ti-E inverted widefield microscope (Perfect Focus System) equipped with a CSU-X1 confocal scanner unit and an Evolve back-illuminated EMCCD camera (Photometrics). Images were recorded using 100x HC Plan APO oil immersion objective (Leica, NA 1.4). Intensities were set to 10% for the 405 nm and 30% or 50% for the 491 nm lasers, with exposure times of 100 ms and 50 ms or 25 ms, respectively. 5 z-stacks with 0.5 *µ*m distances were recorded for each channel. 301 time-lapse images were recorded only in the 491 channel.

#### Live cell imaging of ANCHOR ES cells

Imaging experiments were performed on an inverted Nikon Eclipse Ti microscope equipped with a PFS (perfect focus system), a Yokogawa CSU-X1 confocal spinning disk unit, two sCMOS Photometrics Prime 95B cameras for simultaneous dual acquisition to provide 95% quantum efficiency at 11 *µ*m x 11 *µ*m pixels and a Leica 100x oil objective (HC PL APO 1,4 oil immersion). We excited EGFP and IRFP with a 491 nm (100mw) and a 635-nm laser (>28mW),respectively. We detected green and far red fluorescence with an emission filter using a 525/50 nm and a 708/75 nm detection window, respectively. A thermostated heater (Tokai Hit Stage Top Incubator) allowed for heating at 37*°* C, humidity and CO_2_ control (5%). Time-lapse analysis of GFP and IRFP foci was performed in 2D acquiring 241 time points at a 0.5 s time interval. The system was controlled using Metamorph 7.10 software. Time-lapse was concatenated into single TIFF file.

#### RT-qPCR

RNA was extracted from cells using the Nucleospin RNA extraction kit (Machery-Nagel), then cDNA was prepared with SuperScript IV (Invitrogen), following the manufacturer ‘s instructions and using random hexanucleotides as primers. The cDNA was quantified by qPCR on a LC480 LightCycler (Roche), using QuantitTect SYBR Green PCR kit (Qiagen). Amplification was normalized to GAPDH. Primer sequences are given in Supplemental Table 1.

### Image pre-processing

#### Spot detection and tracking

Spot detection and tracking for all movies was performed with ICY, an image analysis software [36]. Localization precision was then enhanced by assuming that the spots have the shape of a 2D Gaussian function. We optimize its localization using the NM-Simplex method [37] and estimate localization precision using MCMC [38]. This method is implemented and automatically runs while loading trajectories in GP-Tool. For more information see figure S4.

#### Multi-channel alignment correction

For the ANCHOR ES cell line experiments, we used a spinning disk microscope setup with 2 cameras, i.e., one per channel. Even though these cameras were aligned manually using fluorescent beads, we could still observe non-negligible differences between images captured in both cameras. Furthermore, even in rare situations when both cameras were in perfect alignment, we could observe effects of chromatic aberrations towards the edges of the image due the different wavelength used. To correct for such problems, we performed digital post-alignment using a generic set of affine transformations including translation, rotation and scaling as defined in

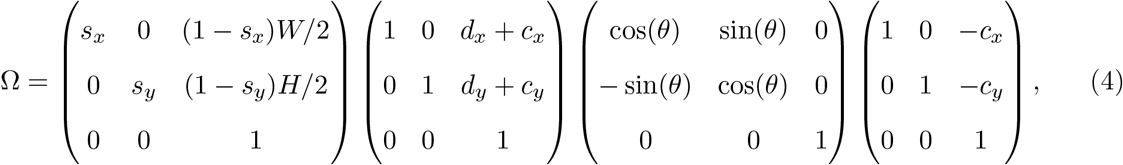

where, *s*_*i*_ accounts for scaling in directions x and y, *d*_*i*_ accounts for translation in both directions and *θ* is the angle of rotation between both channels in relation to point *c*_*i*_.

To infer optimal parameters for correction, we used 10 frames from all the movies recorded in the session and maximize the following likelihood using the Nelder-Mead simplex model [37]

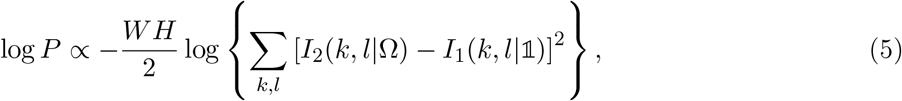

where W and H correspond to width and height of images and *I*_*r*_(*k, l*|*A*) is the value of pixel (k,l) in channel r given transformation A. Fig. S6 shows examples of misaligned images and how the alignment improves greatly after applying our algorithm.

### Derivation of GP-FBM models

#### Covariance function of fractional Brownian Motion

The covariance function of FBM can easily be derived from the assumption of two basic properties: stationary increments *B*(*t*) − *B*(*s*) ∝ *B*(*t* − *s*) and a power-law variance, *B*(*t*)^2^ ∝ |*t*|*α*. Then, the off-diagonal terms of the covariance function can be determined as follows:

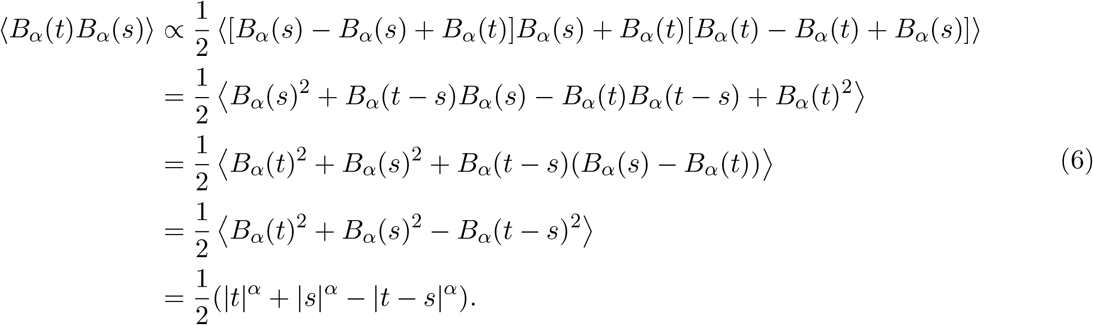

Finally, the apparent diffusion coefficient *D* is introduced as a proportionality factor to re-scale the movement leading to the final kernel as presented in the main text:

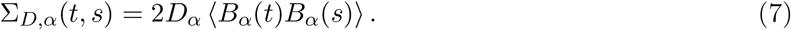

#### Bayesian inference of diffusion parameters

The GP provides the probability of observing a trajectory ***r*** given D and *α*. Then, we applied Bayes theorem [39] to obtain the posterior distribution over the diffusion parameters given the measured trajectory:

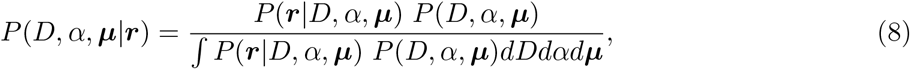

where *P* (*D, α*, ***µ***) represents the prior distribution of the model parameters. Assuming a flat prior on ***µ***, *D* and *α*, the log-posterior can be expressed as

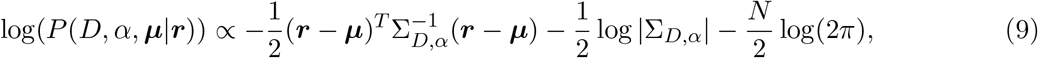

where N represents the number of points measured and | · | is the determinant function. To obtain maximum posterior estimates, we optimized (9) using the Nelder-Mead Simplex method [37]. In addition, to calculate confidence intervals for our estimations we used the MCMC method [38] to sample from the posterior probability distribution. Note that, thanks to this Bayesian approach, available prior knowledge of the diffusion parameters can easily be incorporated into the analysis.

#### Incorporation of substrate movement in the GP-FBM framework

In the main text, we introduced an extended GP-FBM model to deal with external sources of movement that may distort the analysis of particle diffusion. Here, we present the derivation for two particles subject to a common substrate movement, however it can be extended for an arbitrary number of particles using the marginalization rule of multi-variate Gaussian distributions [39]. The key idea is to assume that the movement of the particles with respect to the substrate as well as the movement of the substrate itself can be described by independent fractional Brownian motions. Therefore, the probability of observing the particle trajectories ***a***_**1**_ and ***a***_**2**_ with respect to a given frame of reference that moves with the substrate and the trajectory ***R*** of the moving frame is:

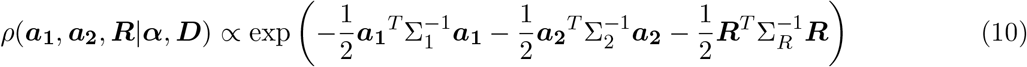

where Σ_1_, Σ_2_ and Σ_*R*_ are FBM covariance matrices that are fully characterized given the diffusion parameters ***D*** = {*D*_1_, *D*_2_, *D*_*R*_} and ***α*** = {*α*_1_, *α*_2_, *α*_*R*_}.

Next, to obtain the probability distribution over the particle trajectories ***r***_**1**_ and ***r***_**2**_ with respect to the microscope reference frame, we applied the the change of coordinates ***r***_*i*_ = ***a***_*i*_ + ***R*** (see scheme in Fig. 2a) leading to the matrix expression,

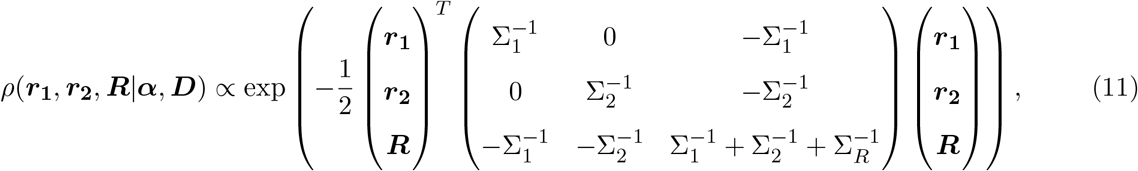

Then, to marginalize out the unobserved trajectory of the moving frame ***R***_*i*_, we need to calculate the inverse of the block matrix in equation 11. To do so, we use results from [40] on inverting a 2×2 block matrix such as

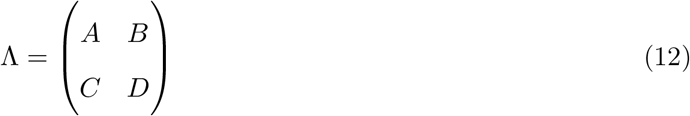

according to the following result

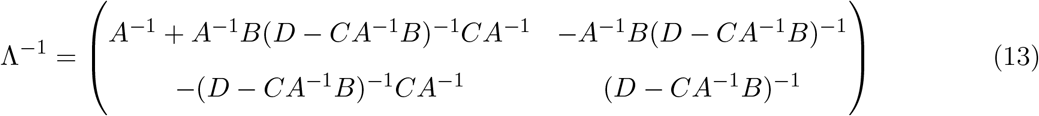

Taking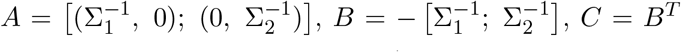, and 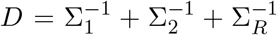, it is easily shown that the top-left corner of Λ^*−*1^ is given by [(Σ_1_ + Σ_*R*_, Σ_*R*_); (Σ_*R*_, Σ_2_ + Σ_*R*_)]. Using this result, we can marginalize out ***R*** in equation 11 giving the expression,

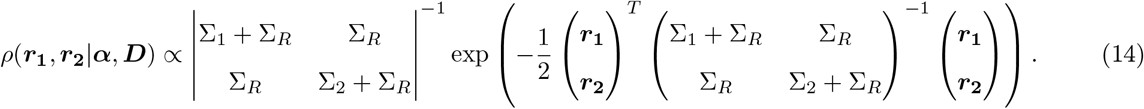

This result clearly shows that the substrate movement induces a correlation between the particle trajectories as the off-diagonal elements of the block matrix in (14) are non zero. In addition, the covariance matrix of the substrate movement appears also in the diagonal terms, increasing the overall variance of the particles and their total movement. Therefore, if this correction is ignored the diffusion parameters are over-estimated.

#### Inference of substrate movement

Similarly as before, the diffusion parameters of the particles as well as the substrate can be estimated using equation 14 and Bayesian inference. Furthermore, we can also estimate how large the movement of the substrate is. For that, we can calculate the conditional distribution of ***R*** given the particle trajectories as

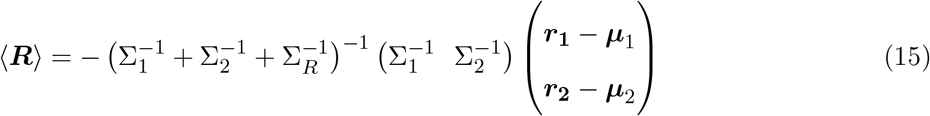

with co-variance matrix given by 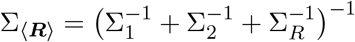.

Unfortunately, we could not find an analytical solution for this problem. Nonetheless, we can solve it numerically. In Fig. S1a we show an example of the estimated movement for the substrate with one standard deviation compared to the real simulated trajectory for a system with two particles. In Fig. S1b-c we display the overall accuracy when working with two or more particles.

### Benchmarking GP-FBM

#### Simulated trajectories

We simulated single trajectories with 250 time points using the aforementioned Gaussian process with FBM kernel. Similarly to experimental setup, trajectories are simulated using time steps of 0.5 seconds and positional noise was included. To benchmark a system of N particles affected by substrate movement, we generate N+1 trajectories and add the latter to all the others. Finally, 10% of points are randomly removed from each trajectory to simulate for experimental occlusions.

#### Mean squared displacement (MSD) implementation

To calculate *D* and *α* for single trajectories, we estimate a MSD using a sliding window method. This method is mathematically defined as follows

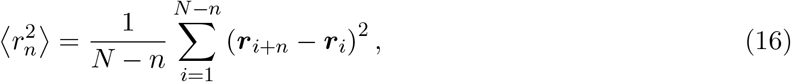

for a trajectory with *N* points and step interval *n*. Due to implicit correlations present in single trajectories, we use only initial 10% step intervals. To improve accuracy, we also estimate an average localization error *σ*. Finally, this experimental curve is approximated by the theoretical mean squared displacement equation

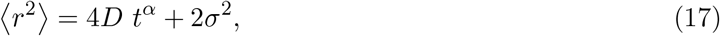

from which diffusion parameters are inferred using linear regression. For more information [41].

#### Displacement distribution based (DDB) implementation

The theoretical expressions for the displacement distribution is obtained as a solution of the Fokker-Planck equation with localization error *σ*. In polar coordinates it takes the form

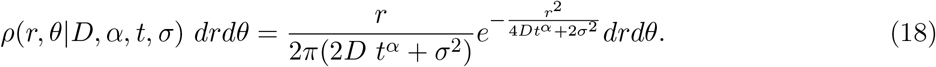

In order to calculate experimental distributions for single cells, we resort to a sliding window method similar to the one present for MSD. Differently, we calculate normalize histograms with all the absolute displacement values. As before, we calculate an average localization error *σ* to improve localization and use only histograms calculate for initial 10 step intervals. With these measurements, we optimize the equation above for D and *α* using Bayes approach with non-informative priors for both parameters.

#### Statistical analysis

To compare diffusive properties or inter-probe distances across different loci or conditions (Figs 3c and d, and 4b-d), we performed Wilcoxon rank sum tests. For inter-probe distances, the distributions of the median distances for each movie were used. For Figs 4c and d, where fifteen pairwise comparisons are possible, the p-values were corrected for multiple testing with the Benjamini-Hochberg method.

#### Use of published epigenomic datasets

ES Hi-C sequence data from [28] were taken from Gene Expression Omnibus (GSE96107), and mapped to mm9 and normalized with FAN-C [42]. The normalized submatrix (chr6:51500000-53000000) was then extracted for visualization. ES and neuronal precursor cell H3K27ac and CTCF ChIP-seq data were taken as bigWig files from Gene Expression Omnibus; GSE96107 for all except ES H3K27ac, taken from GSE49847) and visualized in R using the package *rtracklayer*.

## Acknowledgements

We thank Luca Giorgetti for providing the TetO ES line and for critical reading of the manuscript. This work was possible thanks to funding from grants by LabEx INRT (ANR-10-LABX-0030-INRT, a French State fund managed by the Agence Nationale de la Recherche under the frame program Investissements d ‘Avenir ANR-10-IDEX-0002-02), CNRS «Osez l ‘interdisciplinarité !», ERC (Starting Grant 678624 - CHROMTOPOLOGY) and ATIP-Avenir. The microscopy was performed at the Imaging Center of the IGBMC.

## Supplementary Information

**Table S1:**
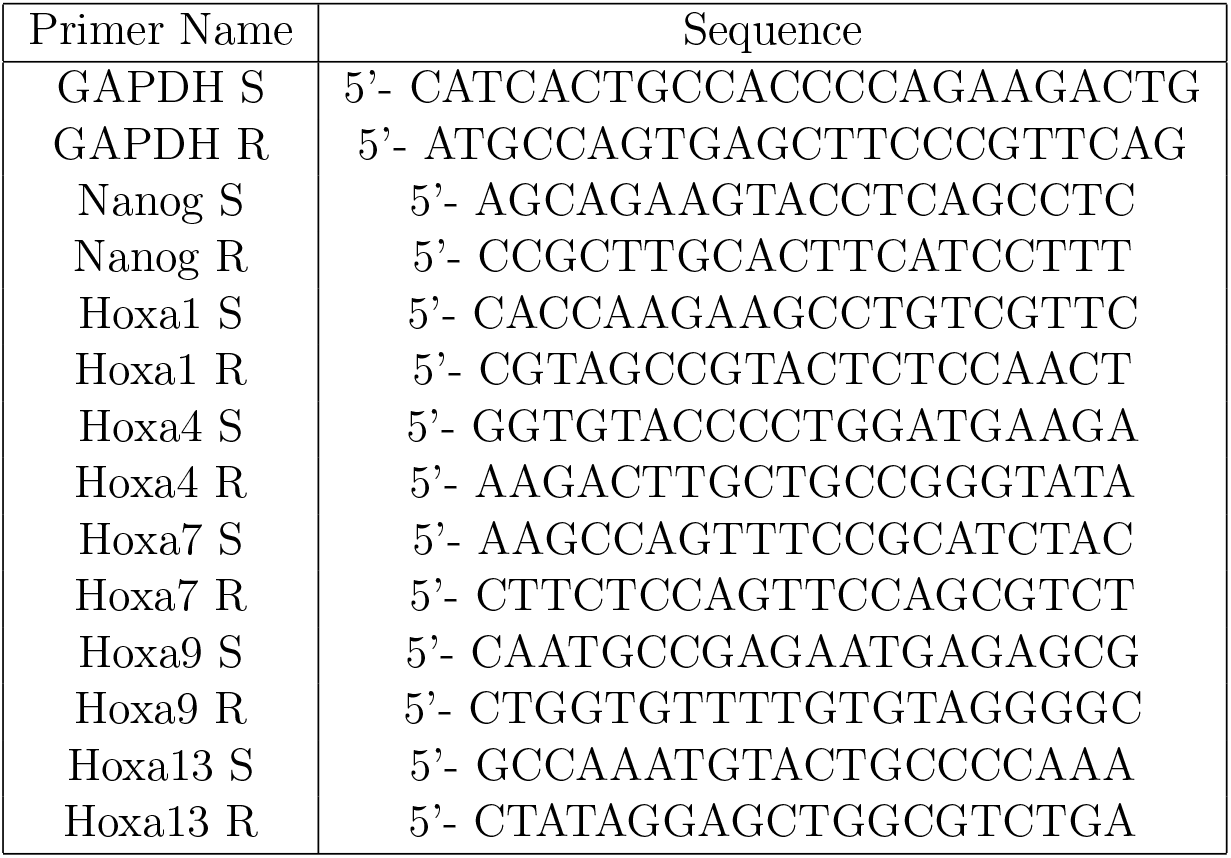
Primers

**Figure S1:**
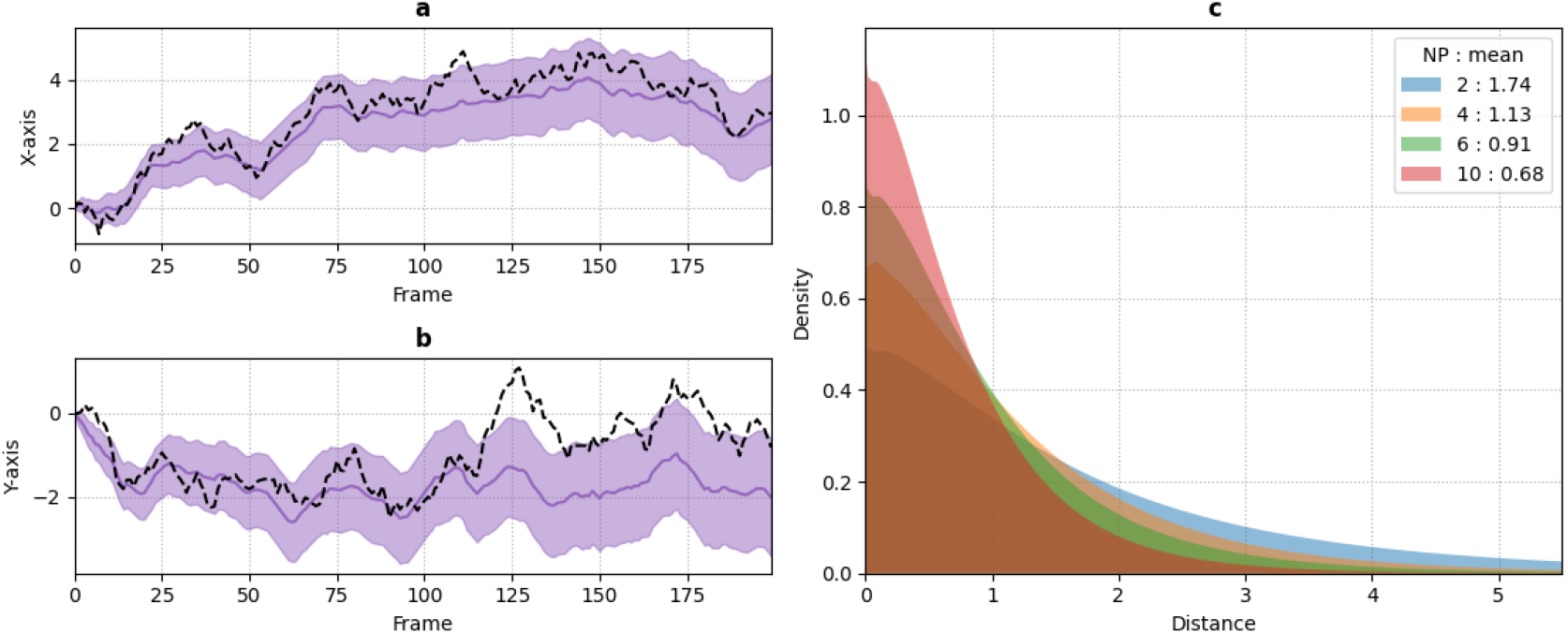
Estimating background movement from tracked particles. (a) Black dashed lines correspond to simulated substrate movement, while purple line is the path estimated for the substrate using GP-FBM. Shaded area correspond to one standard deviation. (b) Positional error distribution in pixel units calculated between substrate simulated and estimated trajectories for systems with 2, 4, 6 and 10 particles.

**Figure S2:**
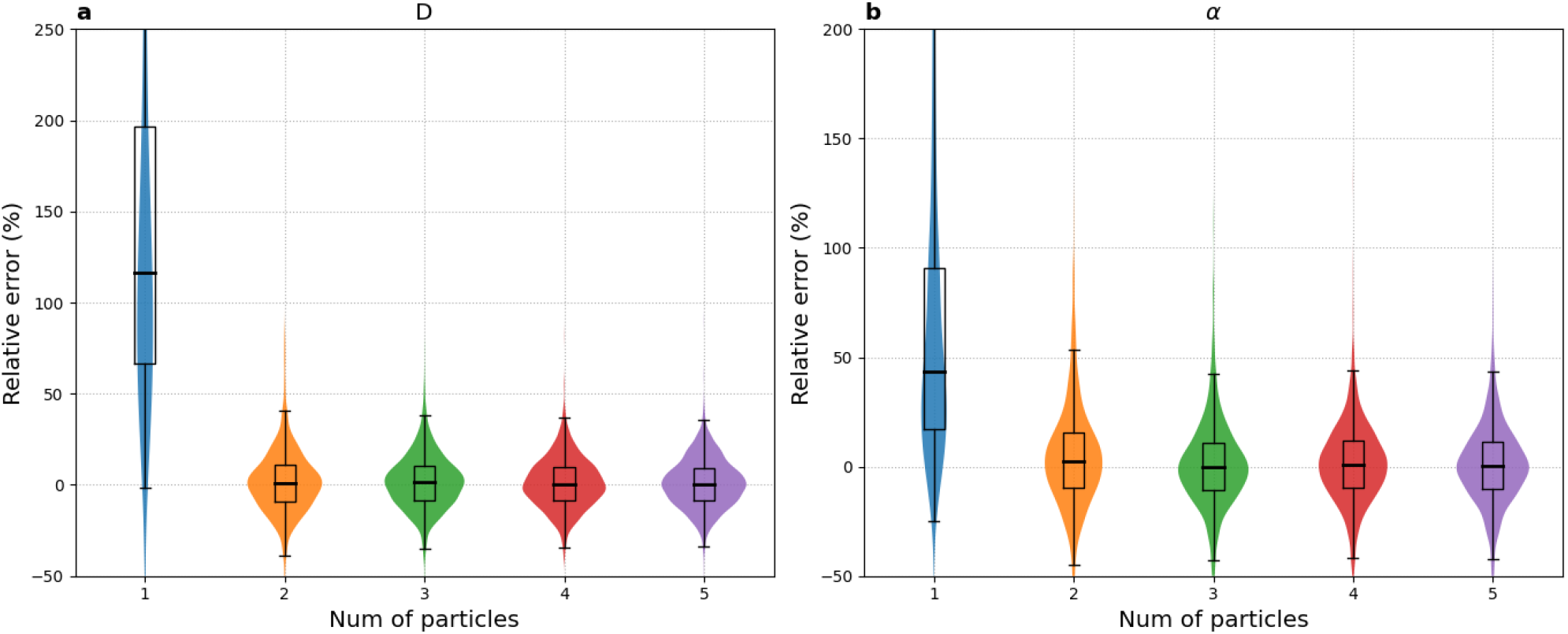
Accuracy gain with number of particles when correcting for substrate movement. If more than one particle under similar context are present, we can used implicit correlations to correct inference. As we can see, there is little gain in precision when using more than 3-4 particles.

**Figure S3:**
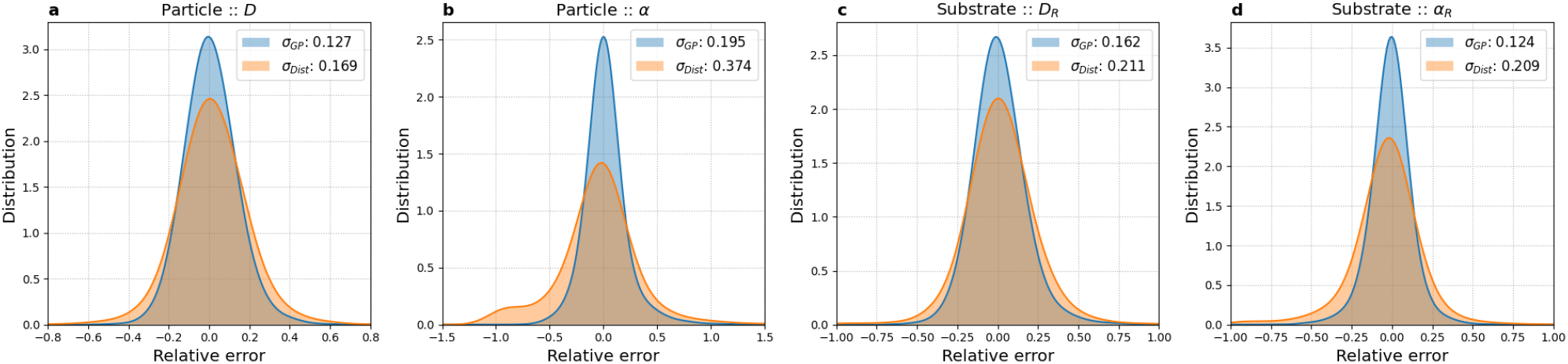
Bench-marking GP-FBM and DDB approaches for inference of diffusion properties with substrate movement. A set of 10,000 trajectories of two particles with substrate movement were generated. Simulated values were inferred using GP-FBM (in blue) and DDB (in orange) and relative errors on *D* and *α* were plotted for particles and substrate. *σ*-values correspond to one standard deviation.

**Figure S4:**
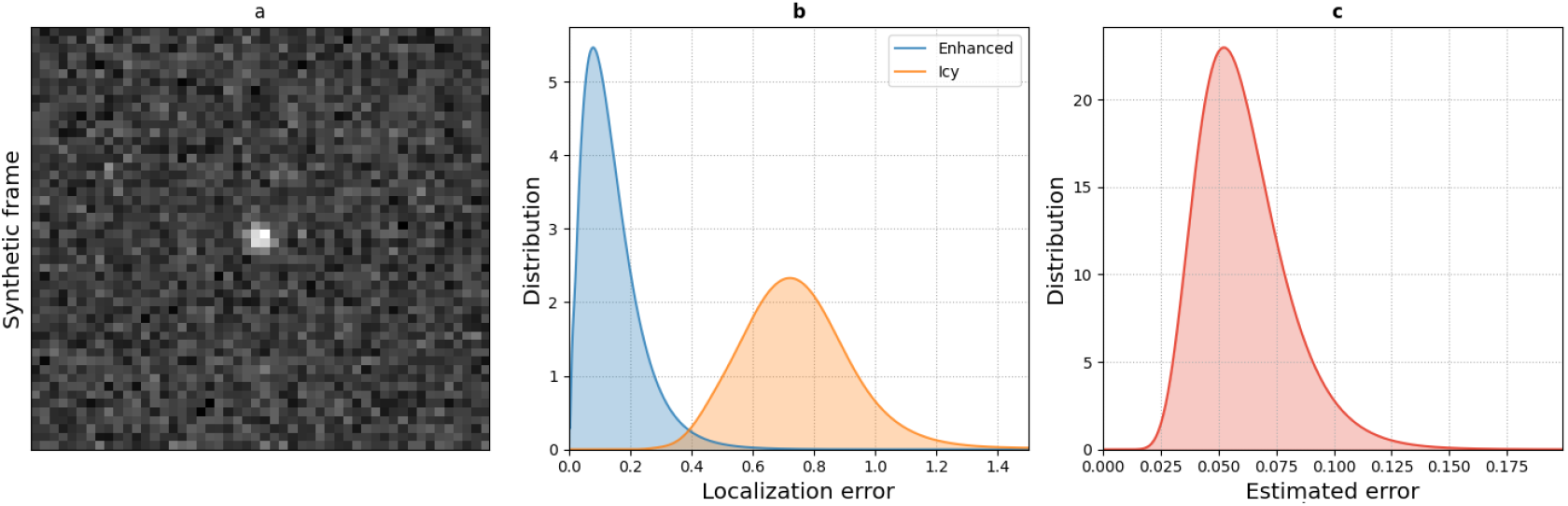
Verifying localization enhancement algorithm via a synthetic movie. To test this method, we generated a synthetic movie with 500 frames for a single particle trajectory. The spot was generated using a 2D symmetric Gaussian shape with deviation 1, background signal was set to 100 and spot intensity to 200. Signal noise was generated in the form of a Poisson distribution by taking original signal as mean. In (a) we have a frame of such a synthetic movie. (b) Spots were tracked with ICY and further enhanced as described in Methods. Distribution of positional errors were calculated using simulated trajectories as reference. (c) Distribution of positional error standard deviations, computed per spot with MCMC.

**Figure S5:**
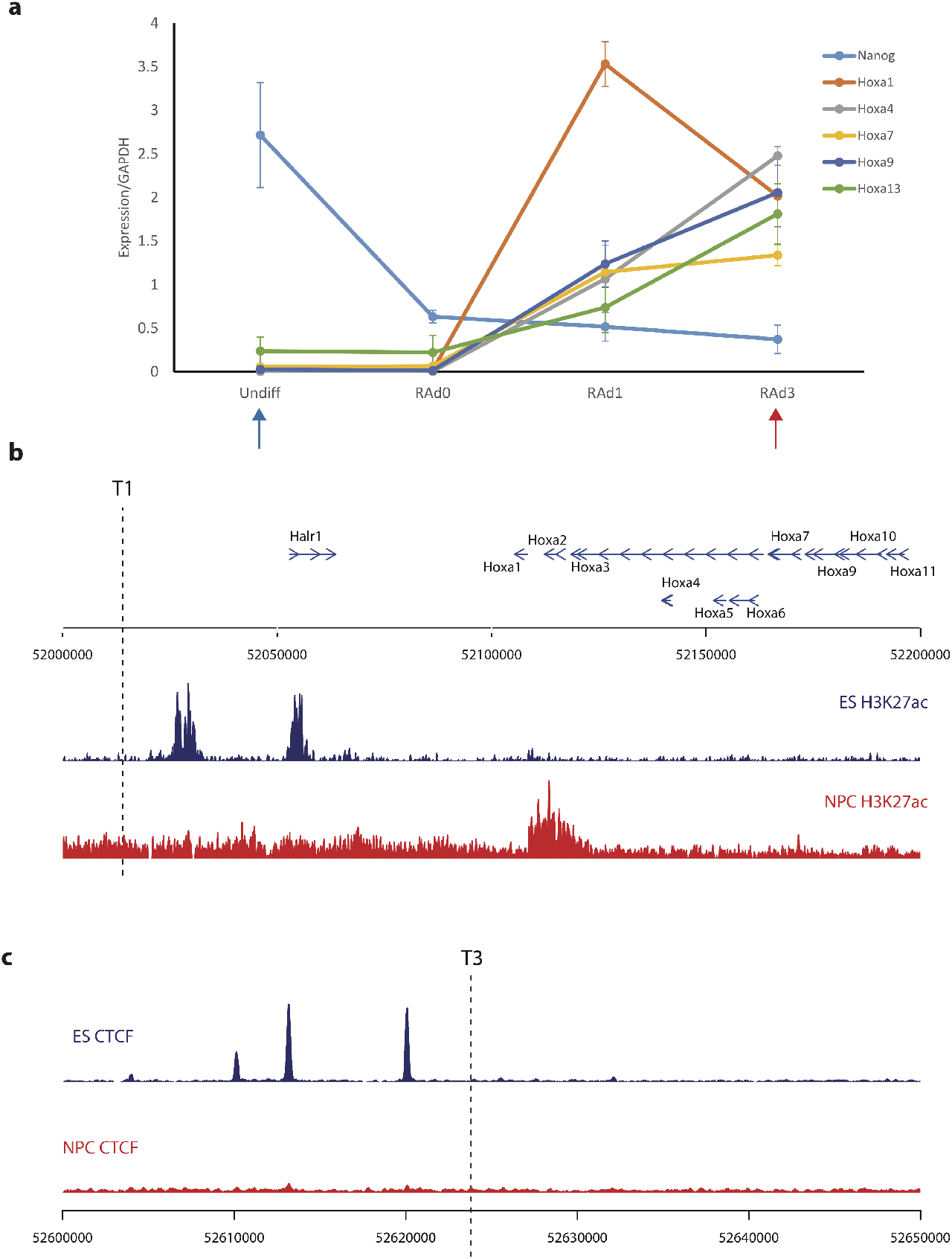
(a) RT-qPCR results, showing expression of Nanog and HoxA genes, normalized to GAPDH, in ES ANCHOR cells before and at different stages of induction by retinoic acid. The graph shows the average results over triplicate experiments. The blue and red arrows indicate the timepoints at which microscopy is performed, as in Fig 4. (b) Gene positions and H3K27ac ChIP-seq tracks in ES (blue) and neuronal precursor cells (red) are plotted for a 200 kb window containing the HoxA cluster and the position of ANCHOR label T1. T1 is <15 kb from an ES-specific putative enhancer. (c) CTCF ChIP-seq tracks in ES (blue) and neuronal precursor cells (red) are plotted for a 50 kb window flanking the ANCHOR label T3. T3 is very close to ES-specific CTCF sites.

**Figure S6:**
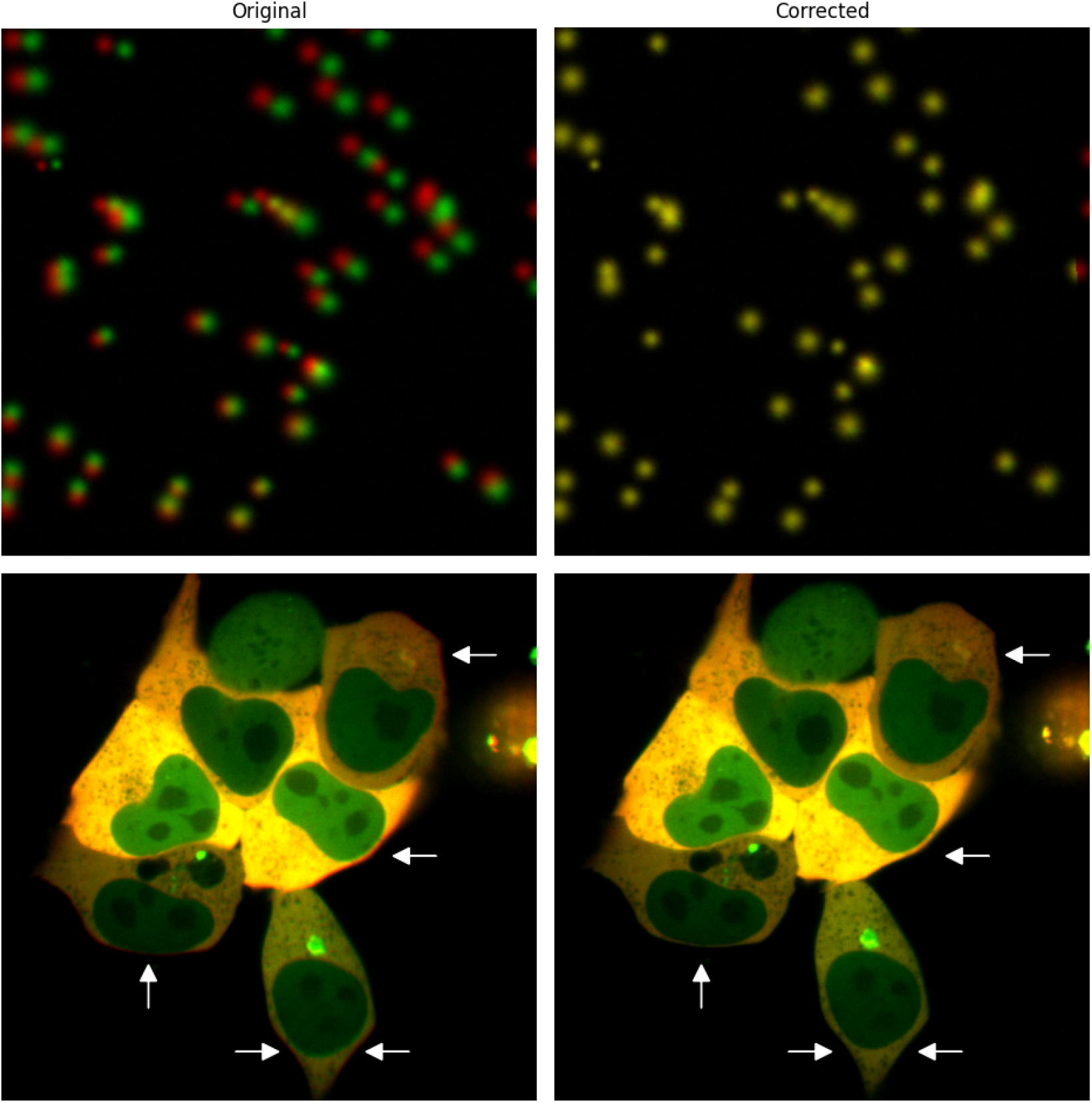
Top images display simulated spots in which we exaggerate effects of camera misalignment. On the bottom, white arrows point to regions with pronounced effects caused by chromatic aberrations. After alignment algorithm is deployed, both channels are properly superposed.

